# Force sensing by Piezo1 regulates endothelial secretory granule exocytosis

**DOI:** 10.1101/2025.09.26.678766

**Authors:** Sammy El-Mansi, Mina Al-Saegh, Loïc Rolas, Matthew Golding, Mathieu-Benoit Voisin, Sussan Nourshargh, Thomas D. Nightingale

**Affiliations:** Centre for Microvascular Research, William Harvey Research Institute, Barts and the London School of Medicine and Dentistry, Queen Mary University of London, London EC1M 6BQ, UK

**Author notes:** **CORRESPONDANCE** Sammy El-Mansi and Thomas D. Nightingale, Centre for Microvascular Research, William Harvey Research Institute, Charterhouse Square, Faculty of Medicine and Dentistry, Queen Mary University of London, London EC1M 6BQ, United Kingdom; and.

## Abstract

Endothelial cells (ECs) rapidly alter their phenotype to support haemostasis and inflammation through the exocytosis of specialised storage organelles, known as Weibel-Palade bodies (WPBs). This exocytic response is stimulated by a variety of physiological and pathological triggers which act through well-described transmembrane receptors. However, the influence of mechanical forces and how they shape this fundamental endothelial response remains relatively unexplored. Here we demonstrate that opening of the mechanosensitive cation channel Piezo1 on the EC surface triggers rapid WPB exocytosis. The dynamics and mechanisms were investigated using the Piezo1 agonist (Yoda1) and by modelling mechanical opening through EC-ICAM-1 ligation by THP-1 cells under low shear flow. *In vitro*, localised von Willebrand Factor (VWF) secretion occurred at THP-1 adhesion sites, suggesting a reciprocity of VWF release alongside leukocyte capture. *In vivo*, Yoda1 stimulation of tissues resulted in a dramatic increase in neutrophil and platelet adhesion to postcapillary venular walls, spatially aligned with cargo release (VWF). The rapid dynamics of this response was revealed by the application of high resolution confocal intravital microscopy to transgenic *EGFP-Rab27a* mice that exhibit fluorescent WPBs and neutrophils. These data reveal a previously unrecognised instigator of WPB exocytosis and uncover a mechanosensitive molecular pathway coupling inflammation with haemostasis.

**GRAPHICAL ABSTRACT:** Graphical abstract: Activation of Piezo1 by pharmacological agonism or leukocyte adherence to ICAM-1 under low flow promotes VWF secretion and platelet capture through a protein kinase C-dependent mechanism.

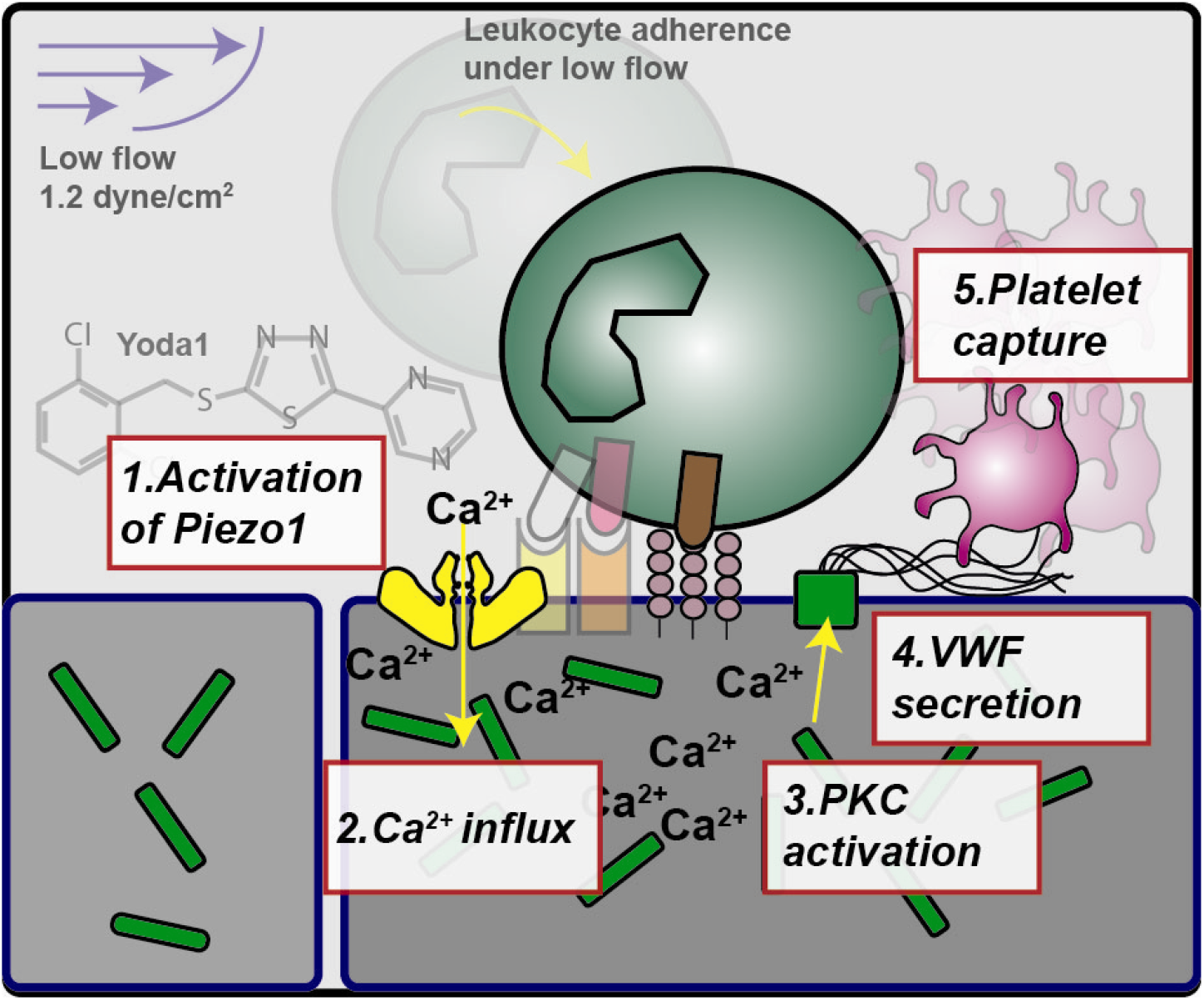

**SUMMARY:** This research describes how the mechanosensitive cation channel Piezo1 elicits the rapid exocytosis of endothelial secretory organelles, provoking platelet capture to sites of leukocyte adherence. The *EGFP-Rab27a* transgenic mouse demonstrates a technical advance, facilitating intravital imaging of endothelial secretory organelles.

## INTRODUCTION

The endothelium is essential for numerous physiological processes including angiogenesis, haemostasis and inflammation. Endothelial cells (ECs) form a semi-permeable barrier allowing gaseous and nutrient exchange restricting leakage of plasma proteins and cells into underlying tissue.^1^ During inflammation, leukocytes traverse this tight barrier to limit tissue infection and/or promote tissue repair through trans-endothelial migration (TEM).^2,3^ This response is tightly regulated, requiring an initial tethering and rolling of leukocytes on the venular endothelium.^3^ Subsequent leukocyte activation promotes their interaction with EC ICAM-1/VCAM-1 stimulating leukocyte crawling and firm adhesion. Attachment of leukocytes to ECs under flow exposes the endothelium to force that is transduced via ICAM-1-cytoskeleton interactions which increases membrane tension and opens the mechanically gated cation channel Piezo1.^4^ This facilitates localised Ca^2+^ influx, a key driver of junctional remodelling,^5-7^ shown to be spatiotemporally coordinated using an endothelial-specific genetically encoded fluorescent calcium reporter, GCaMP3.^8^ In sequence, activation of junctional mechanoreceptor complexes and DAG-activated TRPC6 channels^5,9^ also support immune cell extravasation but the precise temporal order remains unresolved.

Piezo channels do not display homology with any other family of ion channel or receptor.^10^ Their unique structure and the lack of a *bona fide* physiological ligand suggest these channels have evolved for the sole purpose of sensing force. Endothelial Piezo1 is perhaps best known as a fluid shear sensor^11-14^ but studies also report the role of Piezo1 in controlling blood pressure^14^ and junctional opening during leukocyte TEM.^4^

Also important to leukocyte TEM is the exocytosis of endothelial storage organelles Weibel-Palade bodies (WPBs) which contain the leukocyte capturing receptor, P-selectin and its co-regulator CD63.^15^ However, the main constituent of WPB is the haemostatic protein von Willebrand factor (VWF) which is best known for its role in recruiting platelets to the EC surface at sites of injury^16^ but are also recruited to sites of TEM^17,18^ during inflammatory processes in the microvasculature. This response helps maintain vascular barrier function limiting plasma leakage. Paracrine signalling from platelet derived Angiopoietin-1 through endothelial Tie2^19^ reinforces actomyosin-regulated contractile pores.^20^

Given that Ca^2+^ flux is one of the best characterised pathways for WPB exocytosis^21^ and that platelet recruitment is commonly associated with TEM we investigated if there was a direct link between the activation of the force activated cation channel Piezo1 and WPB release. Such localised exocytosis could represent an exquisite molecular mechanism for linking leukocyte diapedesis and platelet recruitment. But despite over a decade of intense Piezo1-research efforts the role for endothelial Piezo1 in WPB exocytosis has not been formally investigated.

Using a variety of pharmacological and mechanobiological *in vitro* assays, we demonstrate that opening of EC-Piezo1 induces spatially restricted endothelial secretory granule exocytosis. Taking advantage of a transgenic *EGFP-Rab27a* mouse that exhibits fluorescent WPBs, the application of 4D intravital microscopy for the first time revealed the dynamics of WPB movement in conjunction with neutrophil, platelet and VWF accumulation in venules. These data highlight an overlooked mechanism of WPB exocytosis which has far reaching implications in the aetiology and therapeutic targeting of thrombotic and thrombo-inflammatory pathologies.

## METHODS

### Animals

The transgenic (RAB27A-EGFP/Rab27a)84Seab^22^ mouse line (*EGFP-Rab27a*) was purchased from Jackson Laboratories and heterozygotes maintained through genetic intercrossing with C57BL/6J mice (Charles River, UK). Male mice (8-10 weeks old) were used in experimental procedures.

All experimental procedures using animals were performed in alignment with the QMUL Animal Welfare Ethical Review Body (AWERB) and The Animals (Scientific Procedures) Act 1986 guidelines from the UK Home Office. Animals were housed under controlled environmental conditions (12-hour light/dark cycles at ambient temperature and humidity) on a standard ad-libitum chow diet.

### Intravital and fixed wholemount imaging endothelial secretory organelles

Anaesthetized (3-5% isoflurane) mice were subjected to intra-scrotal injection of AF555/647 fluorescently labelled non-blocking anti-PECAM-1 monoclonal antibody (4 µg, clone 390) (eBioscience Cat:16-0311-85) to fluorescently label the cremasteric microvascular endothelium as previously described.^23^ Two hours later, mice were placed under terminal anaesthesia (intraperitoneal injection of xylazine (10 mg/kg) and ketamine (100 mg/kg)). Surgical exteriorisation of the cremaster muscle and confocal intravital imaging were performed as previously described^23,24^ using an upright Leica SP8 microscope incorporating a 63x water immersion objective lens. Cremaster tissues were superfused with warm (37C) tyrodes’ solution (Control) alone or with Yoda1 (5 µM) or histamine (30 µM) to stimulate Piezo-1 or to induce vascular permeability, respectively. Post-capillary venules of approximately 20-40 µm in diameter were imaged continuously for at least 10 minutes capturing 1µm interval Z-stacks.

To visualise platelets or VWF in vivo, anti-GP1B-DyLight649 (3 µg, Emfret Analytics Cat: X-649) or anti-VWF-AF555 (5 µg, DAKO Cat: A0082), respectively, were injected intravenously immediately prior to surgery.

Whole-mount immunostaining and imaging of cremasteric post-capillary venules were performed as described elsewhere^25^. At least eight independent venular segments were imaged per mouse (n=3 per group) using a Nikon CSU-W1 SoRa spinning disk microscope.

Confocal images (3D) and video recordings (4D) were analysed using Imaris software (Oxford Instruments). Post-capillary venules were identified through EC morphology of blood vessels and positive PECAM-1 staining. Volume surface renderings for ECs, WPB, platelet, VWF were created using the PECAM-1, GP1B, EGFP-Rab27a or anti-VWF-AF555 channels, respectively. using automated settings. Data is presented as the percentage area of each channel in relation to the venular segment area and at least 8 images were analysed per animal.

### Cell culture

Human Umbilical Vein ECs (HUVEC) (PromoCell, Cat: 12203) from pooled donors were cultured as described elsewhere^26^ and used as the cellular model throughout.

### Immunofluorescence of fixed cultured ECs

Commercial suppliers and other relevant information regarding the antibodies utilised here can be found in Table S1. The methodological approach used is described elsewhere.^27^

Confocal imaging of fixed cells was performed using either the Zeiss LSM 800 and Nikon CSU-W1 SoRa spinning disk microscope with 0.1-0.5 µm interval Z stacks.

### In vitro live cell imaging

HUVECs (0.5x10^5)^ plated into four-chamber glass bottom dishes (CELLview, Griener Bio) were transfected with 5 µg of plasmid DNA by electroporation as described previously^28^. Cells were imaged 24 hours post-transfection. Further information regarding the plasmid DNA used here can be found in Table S2.

For calcium imaging, HUVEC were incubated with 5 µM cell permeant Fluo-4, AM (ThermoFisher, Cat: F14201) for 30 minutes before being washed twice in Hanks’ Balanced Salt Solution (HBSS) containing Ca^2+^. 0.5-1 µm interval Z stacks were obtained continuously for 5-10 minutes according to experimental objective using the Zeiss LSM800. Application of Yoda1 (5 µM) in HBSS was applied to the monolayer after the first Z stack had been acquired. Calcium mobilisation plots were obtained using ImageJ by plotting the mean grey value detected in the Fluo-4,AM channel.

### Preparation of antibody coated beads

25µL of sheep anti-mouse IgG coated paramagnetic dynabeads (11201D) were washed three times using PBS. Beads were then coated in 1 µg of mouse anti-human ICAM-1 antibody or mouse control and incubated for 30 minutes with constant rotations. Beads were then washed three times and resuspended in 1mL to a final concentration of 1x10^7^ beads/mL.

### Flow assays using THP-1 cells and anti-ICAM-1 coated beads

We applied and adapted protocols from Wang and colleagues.^4^ HUVECs (3x10^4)^ were plated into 6 well parallel flow chambers (Ibidi™ µ-Slide VI 0.4). Four hours later, HUVEC were left untreated or stimulated with human Tumour Necrosis Factor alpha (TNF) overnight (16 hours). The next day, HUVEC were washed and 3x10^4^ THP-1 cells (ATCC) or ICAM-1 (or mouse IgG control) -coated beads added per well. Antibody coated beads were prepared as per manufacturers’ recommendations (ie. 25µL of sheep anti-mouse IgG coated paramagnetic dynabeads (11201D) were washed three times using PBS. Beads were then coated in 1 µg of mouse anti-human ICAM-1 antibody or mouse control and incubated for 30 minutes) Following a five-minute incubation period, HBSS containing 10 ng/mL anti-VWF-AF555 antibody was passed over the cells at a rate of 1.2 dyne/cm^2^ using a syringe pump. HUVEC were then washed three times in HBSS and fixed using 4% PFA for 15 minutes (room temperature).

### Assessment of target protein inhibition on VWF secretion using NIR fluorescent dot blot

siRNAs targeting Piezo1 (Cat: L-020870-03), were purchased as SMARTpools from Dharmacon (Horizon Discovery). siRNA oligonucleotides targeting luciferase (LUC) were made to order by Eurofins Genomics (sequence 5’ cgu-acg-cgg-aau-acu-ucg 3’). Near-infrared (NIR)-fluorescent dot blot was performed as described previously.^28^

### HUVEC permeability assay

Fibronectin (Biotinylated) (Cytoskeleton Cat: FNR03-A) was prepared at 5 µg/mL and used to coat glass coverslips placed in 12 well plates. HUVEC were then allowed to grow until confluency over a period of 5 days. On the day of assaying, HUVEC were incubated with DMSO, Yoda1 (5µM) or epinephrine (100 µM). 10 minutes after stimulation culture media was replaced with M199 containing Alexa Fluor™ 488-conjugated streptavidin (Thermofisher, Cat: S32354) and incubated for 5 minutes. HUVEC were then washed twice in PBS before fixation with 4% paraformaldehyde, permeabilization and IF staining through established approaches.

### Statistical analysis

Unless otherwise stated an unpaired students t-test was used to analyse experiments with two groups and a one-way ANOVA with Tukey’s multiple comparison post hoc test used for more than two groups.

## RESULTS

### Pharmacological activation of EC Piezo1 promotes rapid exocytosis of endothelial secretory organelles *in vitro*

To study the involvement of Piezo1 in WPB exocytosis we utilised its pharmacological agonist, Yoda1, this acts as a molecular wedge, allowing channel opening in the absence of membrane tension.^29^ Exposure of HUVECs to increasing concentrations of Yoda1 resulted in a dose-dependent release of VWF into the surrounding culture media (**Fig. 1A**). Yoda1 induced rapid and transient exocytosis - with most fusion events observed within the first 30 seconds (**Fig. 1B**). This temporal profile correlated with the peak of cytosolic Ca^2+^ mobilisation (**Fig. 1C** and **Movie 1**). The ability of Yoda1 to stimulate VWF release was selective to activation of Piezo1 as this effect was abolished in Piezo1-ablated siRNA transfected HUVECs (**Fig. 1D and Fig. 1E**). Pan-PKC inhibitors significantly reduced VWF secretion following agonist-induced Piezo1 channel opening (**Fig. 1F**), findings that are in agreement with previous reports showing PKC-dependent release of VWF.^30,31^,^32,33^ Functionally, as expected, Yoda1 significantly increased junctional permeability (**Fig. 1G and 1H**). Collectively, the data demonstrates that pharmacological activation of EC Piezo1 stimulates exocytosis of WPBs, a response that is rapid and PKC-dependent. This suggests a reciprocal Piezo1-dependent relationship between endothelial permeability and exocytosis.

**Figure 1:**
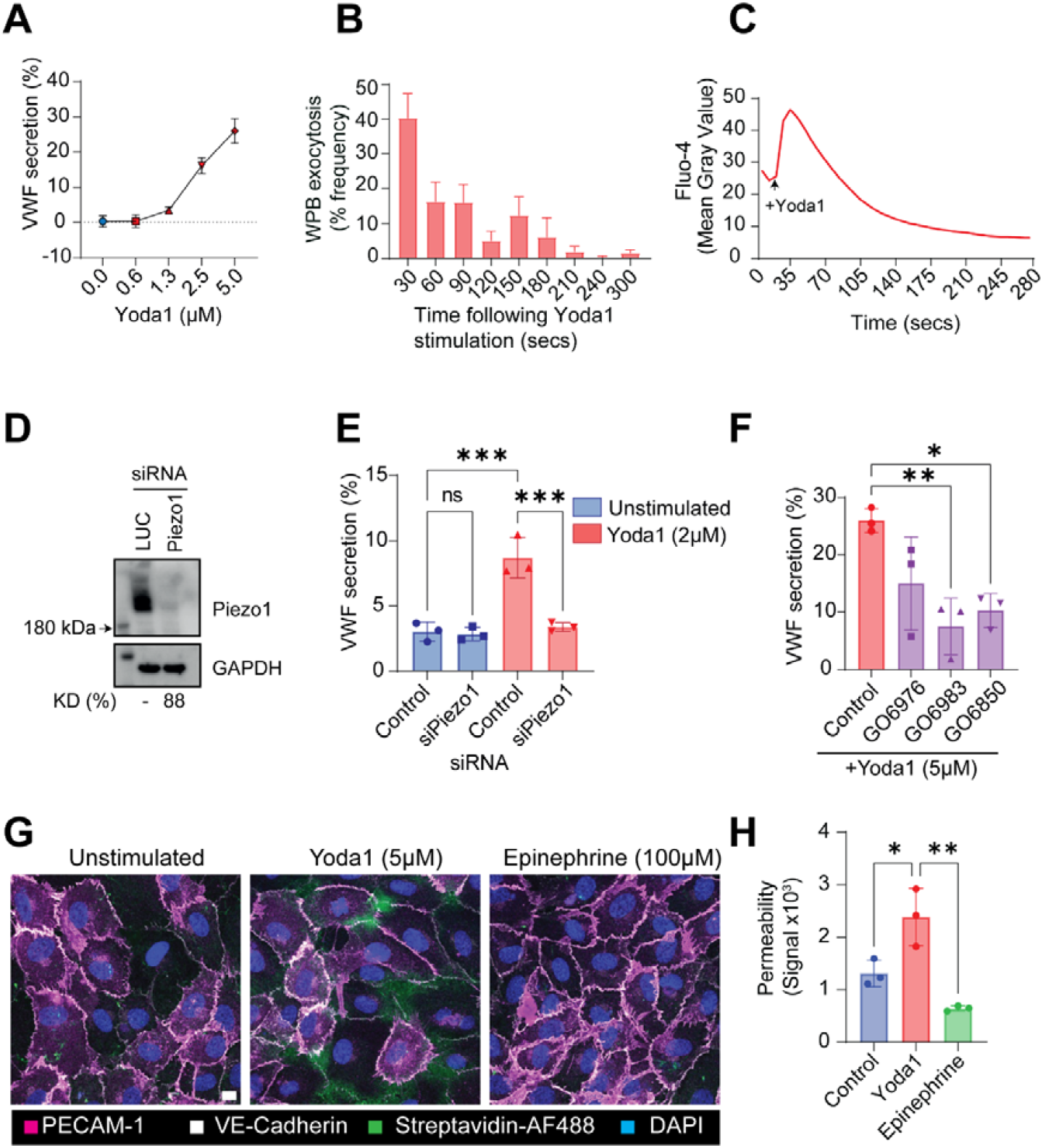
Piezo1 opening triggers rapid exocytosis of secretory organelles alongside junctional permeability. **(A)** Quantification of VWF secretion in response to pharmacological activation of Piezo1. HUVECs were treated with increasing concentrations of Yoda1 before VWF secretion into the culture media was measured using NIR dot blot. Representative data shown. **(B)** Quantification of the temporal dynamics of WPB exocytosis (fusion events) in response to Yoda1 stimulation as determined by live-cell imaging of HUVECs expressing GFP-VWF. A total of 129 fusion events were analysed across 9 cells in 5-minute videos. **(C)** Live-cell imaging of calcium mobilization using Fluo-4-AM dye. Graphical representation of calcium mobilization as mean grey value **(D)** Immunoblot of Piezo1 and GAPDH in HUVECs transfected with 300 pM siRNA targeting firefly luciferase (Control) or Piezo1. **(E)** Yoda1-induced VWF secretion requires Piezo1. N=3 **(F)** Yoda1-induced VWF secretion is dependent on protein kinase C (PKC) isoforms. N=3. **(G)** Confocal imaging to determine EC junctional permeability. Permeability was quantified through the ability of fluorescently conjugated streptavidin (green) to access biotinylated fibronectin below the endothelial cell monolayer. **(H)** Analysis of junctional permeability in response to Yoda1 and Epinephrine. N=3 One way ANOVA with Tukey’s multiple comparison post hoc test. *p<0.05 **p<0.01 ***p<0.005. Control = DMSO vehicle control.

**Figure 2:**
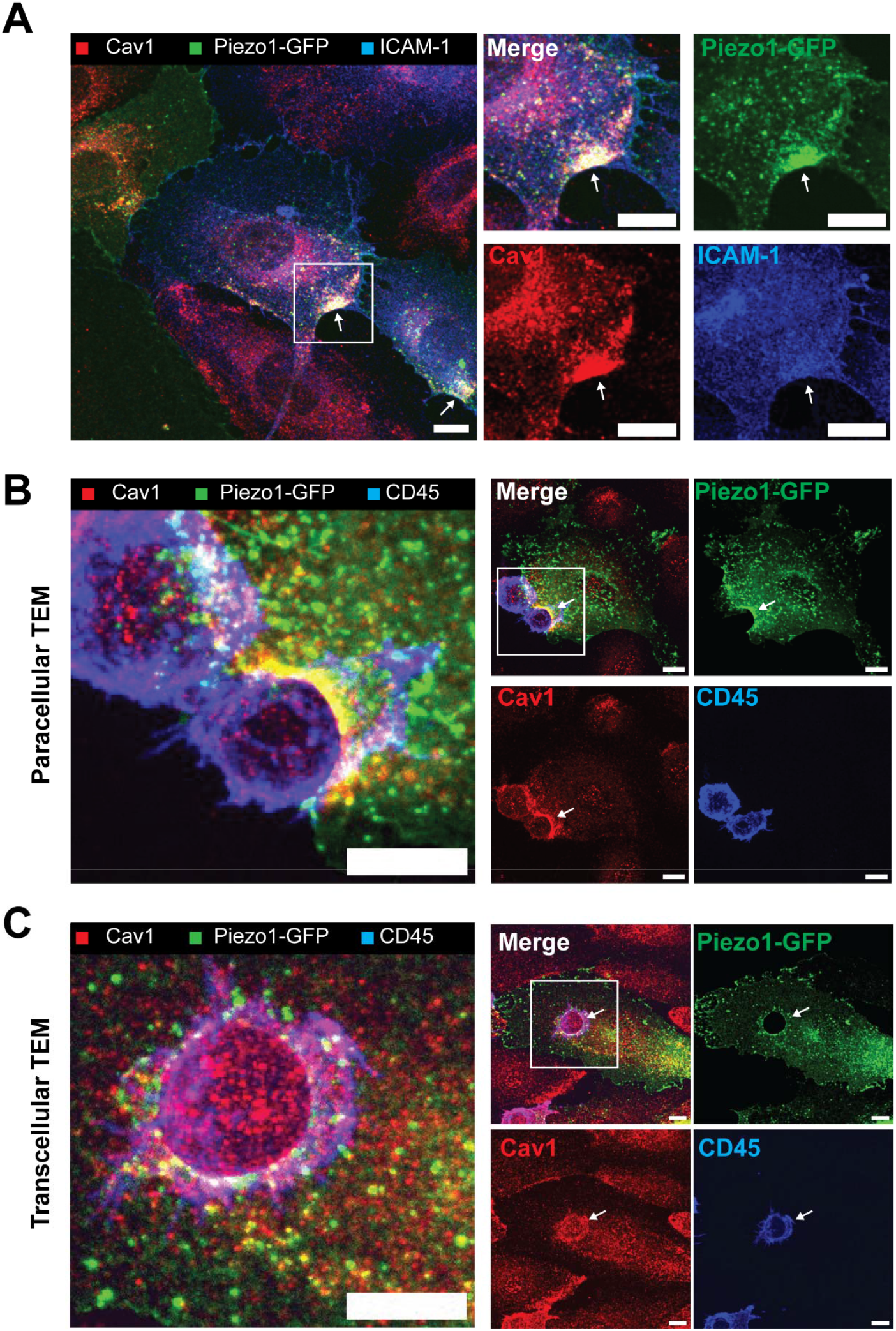
Piezo1-GFP and Caveolin-1 are enriched at the site of leukocyte TEM in vitro. **(A)** HUVEC were transfected with Piezo1-GFP 8 hours before stimulation with TNF (10ng/mL) 16 hours. Immunostaining for Cav1 and ICAM-1 indicated the areas rich in Cav1/ICAM-1/Piezo1-GFP signal. Scale bar 10 µm. **(B)** Under low flow conditions (1.2 dyne/cm^2^) Piezo1-GFP and Cav1 were enriched at the site of THP-1 cell (CD45) paracellular **(C)** transcellular transendothelial migration (TEM). Scale bar 10 µm.

### Localisation of Piezo1-GFP in during leukocyte TEM

Recent work has described a role for Piezo1 activation during leukocyte adhesion as a prerequisite for neutrophil TEM.^4^ Owing to the lack of specific antibodies, the localisation of endothelial Piezo1 during TEM has not yet been documented. To address, we imaged the localisation of Piezo1-GFP in TNF inflamed endothelial cells before also assessing localisation during TEM, including in the context of paracellular and transcellular route.

In static cultures, we noticed Piezo1-GFP localisation appeared in vesicular pools, at the cell junctions and in Caveolin-1 (Cav1) / ICAM-1 rich areas (**Fig.2A**). Cav1 is a marker for caveolae, small cholesterol enriched microdomains that form invaginations in the plasma membrane. They are well documented to play a role in endothelial mechanosensation^34^ and their colocalization with Piezo1 channels has previously been described in alveolar type I cells where they synergise to mediate mechanoresponses to stretch.^35^ Also, directly relevant to our findings are the elegant studies from the Ridley laboratory that described the recruitment of ICAM-1 to caveolae during T cell transcellular migration.^36^

Supporting a role in leukocyte TEM^4^, Cav1 and Piezo1-GFP signal was enriched at the site of paracellular THP-1 cell TEM (**Fig.2B** and **Movie 2**). Further, Piezo1-GFP could readily be observed at the site of transcellular migration (**Fig.2C**). The discovery that Piezo1 activation promotes WPB exocytosis (**Fig.1**) in combination with the enrichment of Piezo1 channels at the site of leukocyte TEM, suggested a possible mechanism for spatial-temporal regulation of WPB exocytosis during inflammatory processes. We next sought to investigate this using microfluidic in vitro models and in vivo imaging techniques.

### Mechanical activation of EC Piezo1 promotes localised exocytosis of VWF

Although mechanical forces such as high (or low oscillatory) shear^12,37^ and hypertensive stress^38^ have been shown to induce WPB exocytosis, this has been classically thought to depend on VEGFR2 signalling. To extend our pharmacological studies to mechanical activation of EC Piezo1 we established a model based on an experimental design developed by Offermann’s research group^4^ (**Fig. 3A**), mimicking the force exerted on ECs via binding of leukocytes to ICAM-1 under low fluid shear stress. Within this assay, it’s important to note that 1.2 dyne/cm^2^ is not sufficient to activate Piezo1 in the absence of other mechanical force placed on ECs.^4,37^ This model incorporates TNF-stimulation of HUVECs - to induce ICAM-1 expression (**Fig. S1A**) - and the addition of leukocytes which under low flow (1.2 dyne/cm^2^) increases membrane tension leading to Piezo1 channel opening.^4^ Of note, TNF stimulation also resulted in an increased expression of Piezo1 (**Fig. S1B&C**).

**Figure 3:**
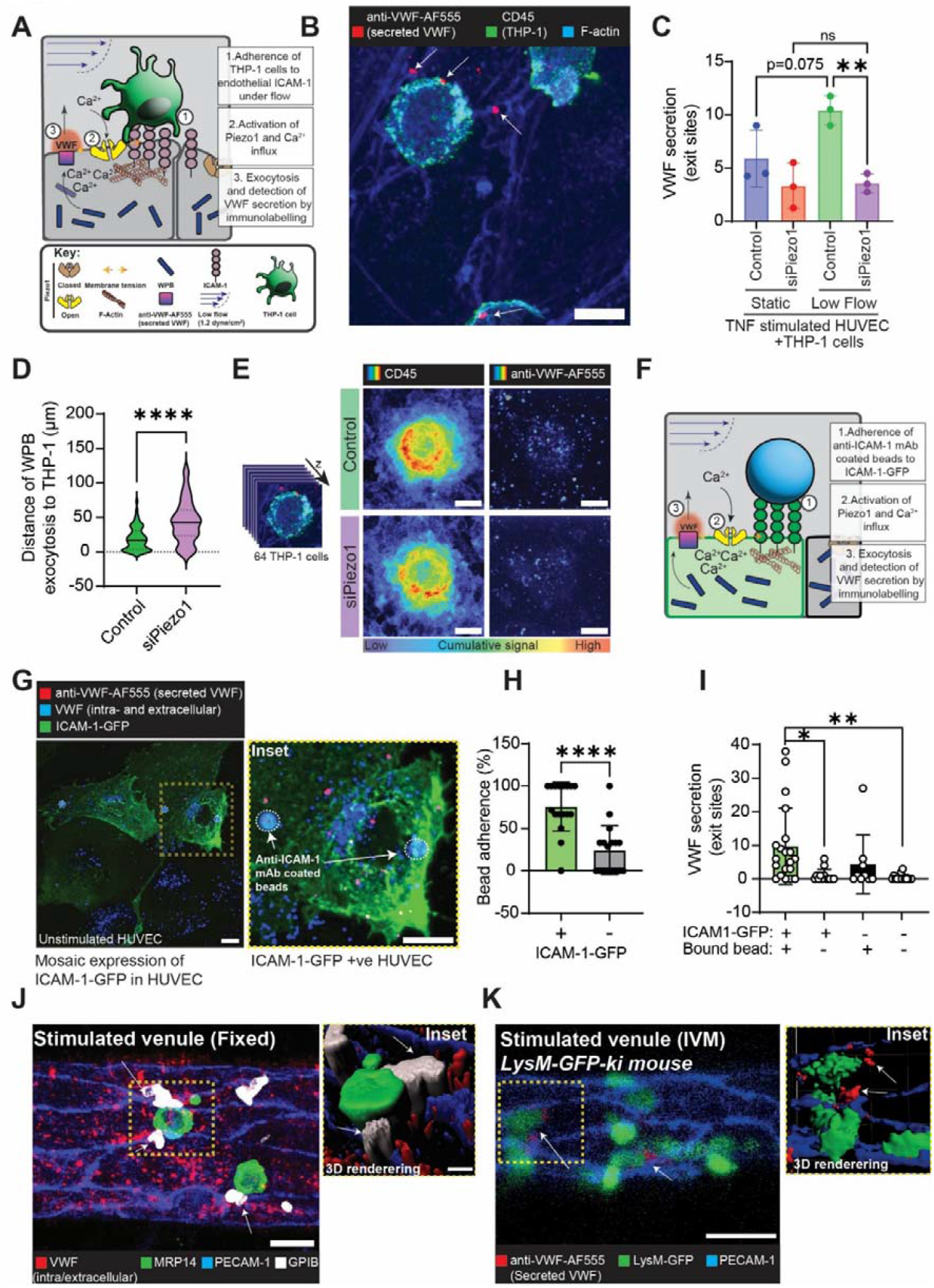
Piezo1 regulates spatial exocytosis of VWF in response to leukocyte adherence under low fluid flow conditions. **(A)** Schematic of in vitro microfluidic set up to study VWF secretion following mechanical activation of Piezo1. Modified from Wang et al., 2022 to detect VWF secretion. **(B)** THP-1 cells were added to either control or Piezo1 knock down HUVEC at a 1:1 ratio under low flow (1.2 dyne/cm^2^). Representative staining shows secreted VWF (red), F-actin (blue), and CD45+ THP-1 cells (green). Scale bar: 10 µm. Arrows indicate secreted VWF at sites of THP-1 adherence. **(C)** The number of puncta representing secreted VWF (exit sites) were quantified under static and low flow conditions. Data is presented as the average number per field of view. n=3 **(D)** Depletion of Piezo1 increased the distance of VWF exit sites from THP-1 cells, indicating that localized VWF secretion is disrupted. **(E)** 400 × 400 pixel insets of 64 different THP-1 cells were concatenated and presented as sum intensity projections. This visually reflects the changes in localized VWF secretion. **(F)** HUVECs were transfected with ICAM-1-GFP to allow anti-ICAM-1 mAb coated bead adherence under low flow (1.2 dyne/cm^2)^ in the absence of inflammatory stimuli. Culture media supplement with anti-VWF-AF555 allowed fluorescent labelling of secreted VWF. **(G)** Confocal imaging of demonstrated mosaic pattern of heterogeneity of ICAM-1-GFP expression and VWF secretion. **(H)** Image analysis determined that anti-ICAM-1 beads bound preferentially to ICAM-1-GFP expressing cells. Students t test. p<0.001 **(I)** Quantification of the VWF secretion in HUVEC positive/negative for ICAM-1-GFP and anti-ICAM-1 binding beads (57 cells from three independent experiments). **(J)** Four-color confocal imaging of wholemount fixed cremaster muscle tissue confirmed platelet adherence at neutrophil adherence sites. **(K)** Anti-VWF-AF555 (5 µg IV) and anti-PECAM-1-AF647 (3 µg intrascrotal) were administered to LysM-GFP mice to visualize VWF secretion during leukocyte adherence in the cremaster muscle. Intravital imaging demonstrated VWF secretion at sites of neutrophil adhesion. Arrows indicate adhered neutrophil encapsulated by VWF signal. Scale bar 30 µm.

Here, an anti-VWF-AF555 antibody was used for extracellular labelling of VWF *in situ* and enabled quantification of the amount and spatial distribution of WPB exocytosis over a short time frame (15 minutes) following addition of THP-1 cells (**Fig. 3B**). This response was most pronounced in low flow, as compared to static conditions and was significantly suppressed following depletion of Piezo1 (**Fig. 3C**). Moreover, under low flow conditions, VWF was secreted in close proximity to sites of adhered THP-1 cells (**Fig 3D**). Cumulative projection of fluorescence intensities of CD45 and extracellular VWF signal illustrates the spatial profile of VWF release at the site of THP-1 cell adherence and its disruption following depletion of Piezo1 (**Fig. 3E**).

Demonstrating whether leukocyte binding to ICAM-1 under low flow promotes WPB exocytosis poses a challenge as ICAM-1 expression requires stimulation by an inflammatory cytokine (TNF), that independently drives some limited but significant, VWF secretion.^16^ To establish a reductionist model, ICAM-1-GFP^39^ was overexpressed in <u>unstimulated</u> HUVECs and anti-ICAM-1 monoclonal antibody (mAb)-coated beads used in lieu of leukocytes. Secreted VWF was again detected through addition of fluorescently conjugated anti-VWF mAb. This created a null system where WPB exocytosis should be minimal unless induced through ICAM-1 ligation and application of force (**Fig. 3F&G**). Anti-ICAM-1 coated beads preferentially bound to ICAM-1-GFP expression HUVEC (**Fig. 3H**). And taking advantage of the mosaic expression profile of ICAM-1-GFP, we noted VWF secretion predominantly in cells exhibiting ICAM-1-GFP expression with adherent beads (**Fig. 3I**).

Having shown that direct activation of Piezo1 in cultured ECs promotes localised VWF secretion and that this can be mimicked by leukocyte adhesion, we sought to investigate this novel secretory mechanism *in vivo*. For this purpose, we analysed TNF-stimulated cremasteric post-capillary venules by confocal microscopy in whole-mount fixed tissue. that were labelled for platelets (anti-GP1B), neutrophils (MRP-4), EC junctions (PECAM-1) and VWF. Here, we observed platelet and neutrophil attachment to junctional regions of venular walls and in close association with VWF (**Fig. 3J**). Building on these findings, we established an intravital microscopy method for real-time detection of VWF secretion in the murine microvasculature. VWF secretion could be detected in vivo through intravenous administration of AF555-anti-VWF antibody to the neutrophil reporter mouse line *LysM-EGFP-ki* in which EC junctions were labelled through local injection of a non-blocking AF647-anti-PECAM-1 antibody.^23,40,41^ In line with the data acquired using our *in vitro* models, this innovative *in vivo* strategy showed secreted VWF in close apposition to adhered neutrophils, indicating spatial restrictions of WPB exocytosis (**Fig. 3K** and **Fig. S2**).

A precedent for localised exocytosis is supported by previous findings - precise areas of increased tension can be induced by deforming the membrane with tether pipettes and this has been demonstrated to induce vesicle fusion in response to Piezo1 activation.^42^ During leukocyte TEM, SNARE fusion machinery (including SNAP23 and syntaxins) are corralled by ICAM-1 clustering^43^ this creates sites of exocytic machinery that favour spatial targeting. By localising VWF release more specifically to sites of TEM the cell mitigates the knock-on effects caused by disruption of the endothelial barrier. We and others^44^ demonstrate that Yoda1 promotes an EC permeability response. Supportive of the findings here, Piezo1 has been shown to be enriched at the cell junctions in the vasculature^45^ and junctions are key sites of leukocyte TEM.^23^ A reciprocal relationship between permeability and exocytosis of VWF induced by local changes in cellular tension allows recruitment of platelets and provides a simple yet exquisitely adapted mechanism to prevent vascular leakage.^19^

### Intravital imaging of endothelial secretory organelles

Since their discovery by electron microscopy (EM) in 1964,^46^ imaging technologies have been paramount to extend our understanding of WPBs. EM revealed their rod-shaped morphology and the subsequent development of GFP-VWF DNA constructs^47^ enabled pioneering live cell imaging of WPB dynamics in vitro. Subsequently, high resolution microscopy and live cell imaging have been used to provide key insights into their biogenesis,^48^ trafficking^49,50^ and regulated exocytosis in vitro.^31,51-53^ However, such findings are limited to cultured endothelial cells and lack the full complexity of the vascular microenvironment in vivo. Here we utilise intravital imaging in combination with an *EGFP-Rab27a* mouse to image WPBs within the microvasculature, capturing their dynamics in the context of blood flow, shear stress, cell-cell interaction and inflammatory signalling. This model represents a step-change in our ability to dissect physiological regulation of WPB trafficking and exocytosis in vivo, with broader implications and applications in homeostasis, inflammation and thrombosis.

The model, originally generated and characterised by Tolmachova and colleagues; uses a transgenic mouse line expressing an EGFP tagged version of Rab27a under the endogenous promotor.^22^ Rab27a is a small GTPase that is resident on the surface of WPBs^54^ but is also recruited to a wide range of secretory granules in a variety of cell types, including neutrophils.^22^ The *EGFP-Rab27a* transgenic mouse therefore exhibits fluorescent WPBs and neutrophils allowing simultaneous comparative monitoring.

Imaging of wholemount fixed cremaster muscles of *EGFP-Rab27a* mice confirmed the presence of EGFP-Rab27a (green) on VWF (red) positive structures within post-capillary venular ECs (PECAM-1-White) (**Fig. 4A**). Furthermore, by comparing Rab27a and VWF labelling we can easily distinguish between secreted and intracellular VWF observed at the point of neutrophil adhesion (**Fig. S3**).

**Figure 4:**
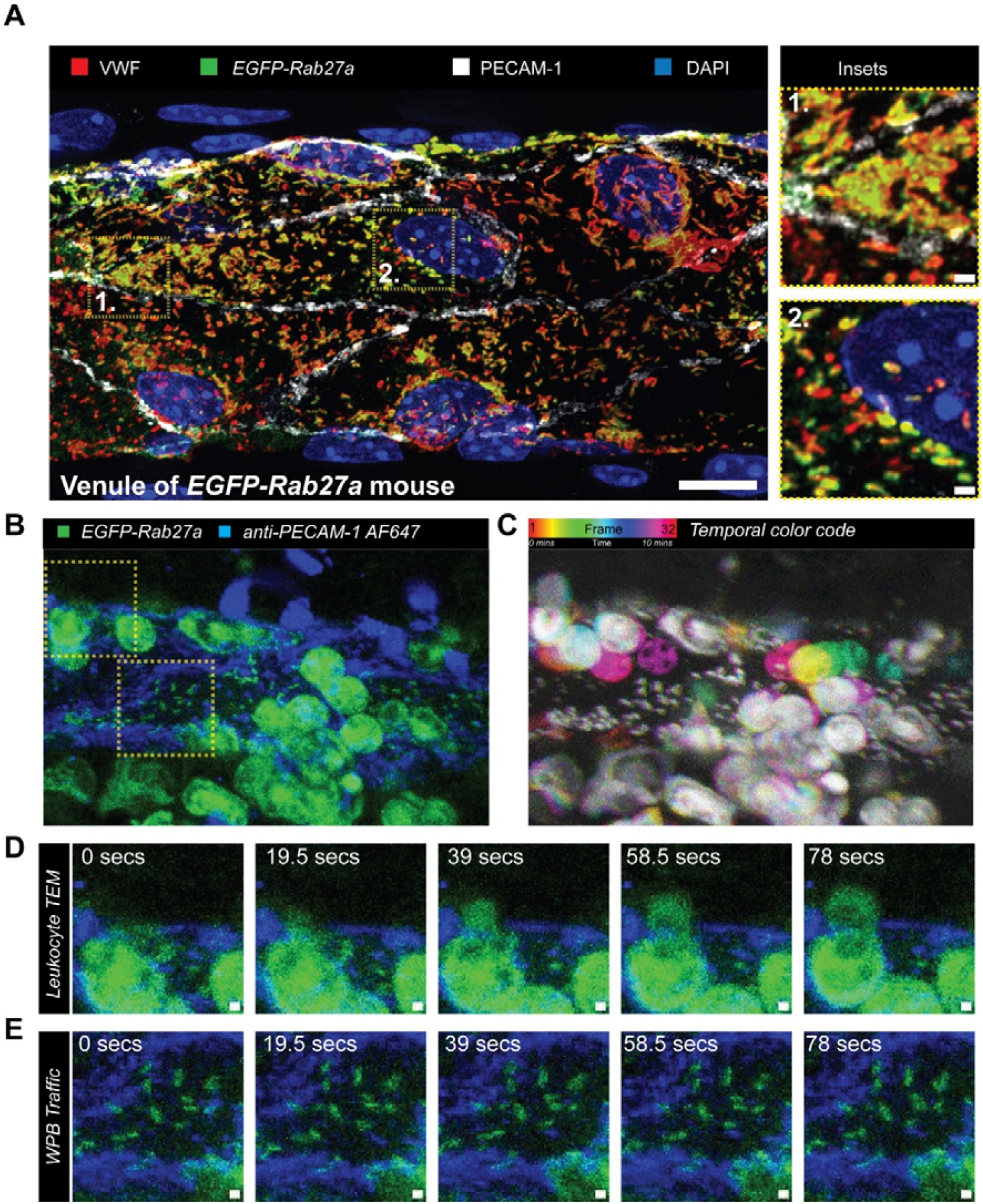
The *EGFP-Rab27a* mouse allows visualisation of WPB dynamics in real time. **(A)** Wholemount fixed tissue confocal imaging of the cremasteric microvascular endothelial cells shows VWF structures (WPBs) positive for EGFP-Rab27a. Scale bar is 10 µm, Inset scale bar 1 µm. Insets show WPBs at 1. tricellular junction and 2. perinuclear region. **(B)** Still from intravital imaging movie. **(C)** Temporal colour coded projection of 10-minute intravital imaging video from histamine stimulated cremaster microvasculature. **(D)** Example of leukocyte extravasation. **(E)** Inset shows WPB dynamics are largely stationary. Scale bar 1 µm.

Using established intravital microscopy protocols^23,55^, including *in vivo* labelling of EC junctions using an anti-PECAM-1 mAb as detailed above, we extended our analysis to real-time imaging of WPB dynamics. These studies represent the first intravital imaging of endothelial secretory organelles (**Fig. 4B** and **Movie 3**). Additionally, the presence of EGFP-Rab27a neutrophils allows the simultaneous monitoring of leukocyte trafficking (**Fig. 4D**).

In contrast to WPBs in cultured ECs that are highly mobile, very little motility of WPBs was observed *in vivo* perhaps indicative of their anchoring on actin structures (**Fig 4C&E**).^49^ However, in a few cases following topical application of histamine (30 µM), long range movement was observed (**Movie 4**). Intravenous administration of fluorophore anti-VWF antibodies was used label to unambiguously label VWF as it is secreted apically into the vessel lumen. In contrast to cells in culture we noted very specific regions of release localised at EC junctions and (in agreement with a role for piezo in VWF secretion) at sites of neutrophil adhesion (**Movie 5 and Fig. S4**). The development of such a model represents a major advance in the field, it enables direct visualization of WPB dynamics in their native vascular environment, under shear and surrounded by all the interacting cell types. This overcomes many of the limitations of in vitro models.

### Pharmacological activation of Piezo1 elicits the exocytosis of endothelial secretory organelles *in vivo*

To test the effect of Piezo1 activation, Yoda1 (5 µM) was applied topically to exteriorised cremaster muscle tissue of anaesthetised *EGFP-Rab27a* mice. This resulted in significant increases in VWF secretion and neutrophil adhesion (**Fig. 5A-C**). Additionally, Yoda1 induced platelet capture to the EC surface (**Fig. 5D&E**), a response that correlated with a reduction in the percentage area of EGFP-Rab27a-positive WPBs (45±3.8% vs 33.4±3.7% p<0.05) (**Fig. 5F**). Finally, we tested whether systemic exposure to Yoda1 could elevate circulating levels of plasma VWF in wild type mice (C57Bl6j). Subcutaneous administration of Yoda1 (2mg/kg^-1^) resulted in a significant increase in plasma VWF levels as compared to control mice (fold increase of 2.2±0.3, p<0.005) (**Fig. 5G**). These findings that are comparable to that observed following subcutaneous epinephrine injection^56,57^ confirming that activation of Piezo1 is sufficient to elicit endothelial secretory organelle exocytosis.

**Figure 5:**
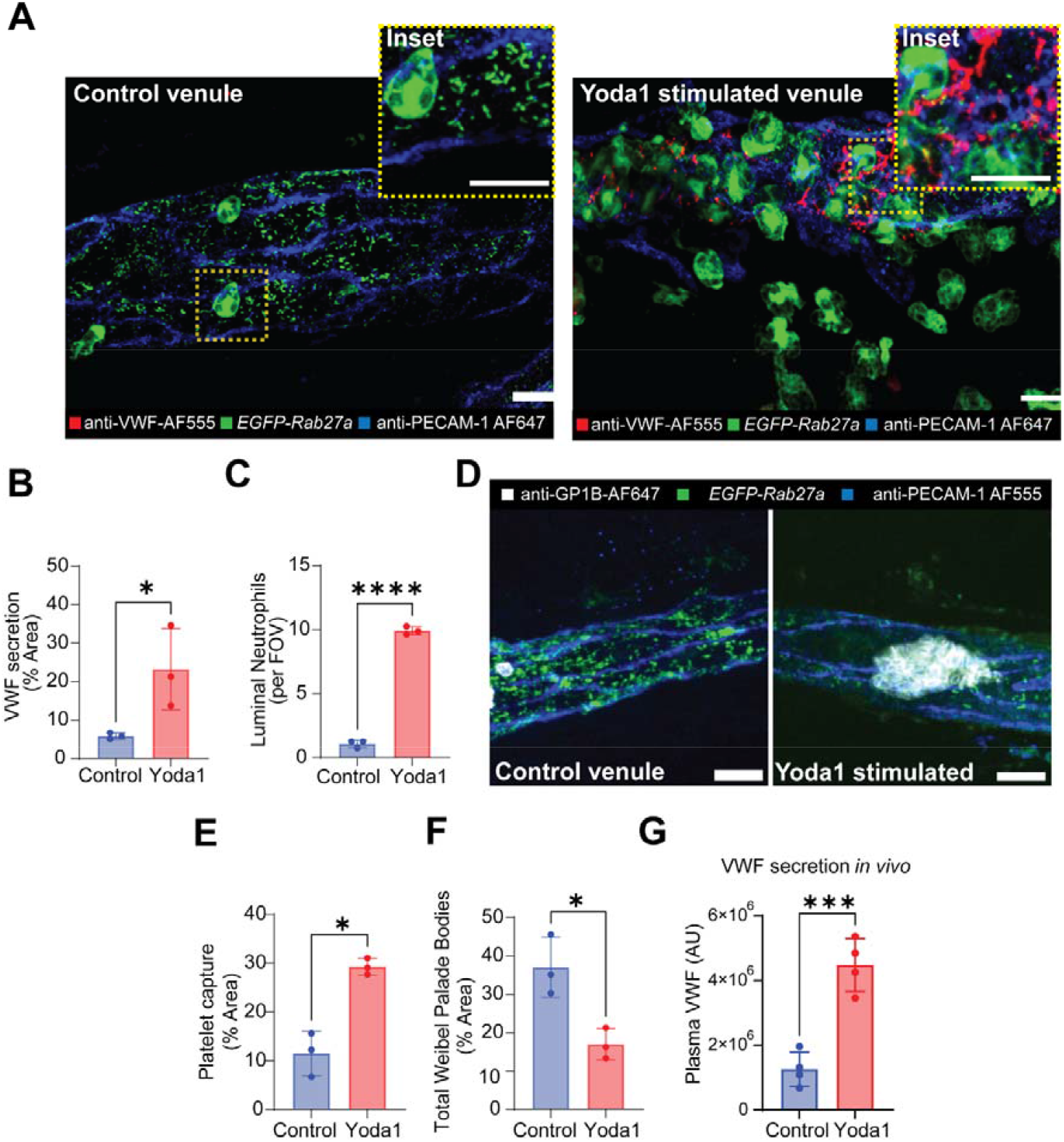
Piezo1 activation promotes VWF secretion *in vivo*. **(A)** In vivo fluorescent labelling of secreted VWF and confocal imaging of post capillary venules from DMSO and Yoda1 treated *EGFP-Rab27a* mice. Scale bar is 10 µm. **(B)** Imaris (Oxford Instruments) was used to quantify the amount of anti-VWF-AF555 signal and normalized to the total area of the blood vessel using PECAM-1 staining as a template. Yoda1 stimulated increased VWF secretion. **(C)** The number of the neutrophils within blood vessel was counted manually. **(D)** Platelets were labelled in vivo through administration of anti-GP1B Dylight649. Quantification of the total area of **(E)** platelets **(F)** EGFP-Rab27a positive WPBs normalized to the vessel area. The number of EGFP-Rab27a WPBs decreased following Yoda1 stimulation. Scale bar is 10 µm. **(G)** Subcutaneous administration of Yoda1 (2mg/kg) increases the amount of VWF in murine (C57Bl/6J) plasma. N=4. Data are mean±SEM from 3 or 4 mice. 8 images from different post-capillary venules (20-40 µm diameter). Students *t* test used throughout. *p<0.05 ***p<0.005 ****p<0.0001.

Dysregulated endothelial Piezo1 activation is likely relevant in a number of pathological contexts^14^. Piezo1 activation occurs in response to high shear^13^ and disturbed flow^37^, EC dysfunction in the aortic arch and the areas surrounding venous valves may be exacerbated by an increase in WPB release. Indeed, VWF deficient mice exhibit delayed progression of atherosclerosis and vascular remodelling (neointima formation) in response to acute vascular injury.^37,58-60^ VWF is also essential in preclinical models of DVT^61^ thus, our findings are relevant to both pathophysiology’s. Our data may also support new approaches to improve the outcomes of coronary artery bypass grafts (CABG). During CABG patient’s autologous saphenous vein are grafted into the arterial vasculature of the heart. This exposes these vessels to high shear rates and often results in either thrombosis (early vein graft failure) or adverse pathological remodelling described as late vein graft failure – a narrowing linked to neointimal hyperplasia and inflammation.^62^ Targeting endothelial Piezo1 channels to limit VWF release and endothelial activation could improve patency.

Collectively, these data demonstrate that activation of Piezo1 drives exocytosis of endothelial secretory organelles in stimulated venules, and by using the *EGFP-Rab27a* mouse line, provide the first real-time imaging of WPB dynamics *in vivo*. Our findings provide an important step forward for the field and a new molecular mechanism underpinning endothelial secretory organelle exocytosis. Further studies will explore the conditions under which Piezo1 contributes to WPB exocytosis, its pathophysiological relevance and its potential as a druggable target to mitigate endothelial dysfunction and thrombo-inflammatory diseases.

## ACKNOWLEDGMENTS

The authors thank their funders and collaborators. They also thank Prof. Manfred Frick and Dr. Giorgio Fois for their guidance related to exocytosis, calcium signalling and imaging. This work was supported by the British Heart Foundation (Grant PG/22/11208 to **T.D.N**, Grant PG/19/73/34663 to **M.V**) and the Wellcome Trust (221699/Z/20/Z to **S.N**). We acknowledge the CMR Advanced Bio-Imaging Facility of QMUL for the help and advice with microscopy.

## AUTHORSHIP CONTRIBUTIONS

**S.E.-M** conceptualised the project, developed the methodology, performed the investigation and wrote the original draft.

**M.A.-S** helped perform the pharmacological studies.

**M.G** and **L.R** provided guidance and technical support.

**M.V** trained and supervised intravital imaging models.

**S.N** provided expert advice and tools for the study.

**T.D.N**. developed the methodology, acquired funding and supervised the study. All authors reviewed and edited the manuscript.

## DISCLOSURE OF CONFLICTS OF INTEREST

The authors declare no competing financial interests.

## Notes

### Competing Interest Statement

The authors have declared no competing interest.

