## Supplementary Materials for "Force sensing by Piezo1 regulates endothelial secretory granule exocytosis"

**This PDF file includes:**

Legends for Movies 1-5.

Tables S1

Figures S1-S4.

**Other Supplementary Materials for this manuscript include the following:**

Movies S1 to S3

**Movie 1: Calcium ion mobilisation in Fluo-4 loaded HUVEC following stimulation with Yoda1 (5 µM).** Movie shows confocal live cell imaging over a 5 minute period (continuous imaging of 0.5 µm interval Z stacks). HUVEC were stimulated with Yoda1 after 5 seconds. Scale bar 10 µm.

**Movie 2: Three-dimensional reconstruction showing Piezo1-GFP localisation at the site of THP-1 cell transendothelial migration.** THP-1 cells were added to TNF inflamed endothelial cells (expressing Piezo1-GFP) before low flow (1.2 dyne/cm2) was applied for 10 minutes. The cocultures were then fixed, immunostained and imaged using spinning disk microscopy. IMARIS was used to create a three-dimensional rendering to illustrate the localisation of Piezo1-GFP during TEM.

**Movie 3: Four-dimensional intravital confocal imaging of WPB dynamics, VWF secretion and neutrophil transendothelial migration (TEM) in a cremaster muscle venule.** This confocal IVM movie shows a cremaster muscle post-capillary venule during of a *EGFP-Rab27a* mouse exhibiting EGFP Weibel-Palade bodies and EGFP neutrophils. EC junctions were stained *in vivo* with a non-blocking AF647-anti-PECAM-1 mAb (blue). Secreted VWF was fluorescently labelled in vivo using AF555-anti-VWF (red). Histamine stimulation (30 µM). The video shows WPBs possibly anchored on actin structures. Continuous imaging of 1 µm interval Z stacks over 10 min time frame.

**Movie 4: Intravital imaging of VWF secretion, WPB dynamics and leukocyte dynamics in histamine stimulated murine cremaster microvascular venules.** This confocal IVM movie shows a cremaster muscle post-capillary venule following exposure to histamine (30 µM) in *EGFP-Rab27a* mouse exhibiting EGFP Weibel-Palade bodies and EGFP neutrophils. EC junctions were stained *in vivo* with a non-blocking AF647-anti-PECAM-1 mAb (blue). Secreted VWF was fluorescently labelled in vivo using AF555-anti-VWF (red). The movie shows exocytosis and long range movement of WPBs possibly via microtubules. Secreted VWF appears junctional and interplay between VWF positive region and neutrophils are observed. . Continuous imaging of 1 µm interval Z stacks over 25 min time frame.

**Movies 5: Three dimensional rendering illustrating the localisation of VWF secretion in the microvasculature.** This movie shows a cremaster muscle post-capillary venule following exposure to histamine (30uM) in *EGFP-Rab27a* mouse exhibiting EGFP Weibel-Palade bodies and EGFP neutrophils. EC junctions were stained *in vivo* with a non-blocking AF647-anti-PECAM-1 mAb (blue). Secreted VWF was fluorescently labelled in vivo using AF555-anti-VWF (red). The movie shows secreted VWF at the tricellular junctions and in close proximity to neutrophils.

| **Target name** | **Host species** | **Cat number** | **Supplier** | **Dilution** | **Method** |
| --- | --- | --- | --- | --- | --- |
| VWF | Rabbit | A0082 | Dako | 1 in 10000 | IF |
| VWF | Sheep | AHP062 | Bio-Rad | 1 in 10000 | IF |
| ICAM-1 | Mouse | sc-8439 | SantaCruz | 1 in 1000 | IF |
| ICAM-1 | Mouse | BBA4 | RandD | see methods | Beads |
| Tubulin | Mouse | T5201 | Sigma Aldrich | 1 in 3000 | WB |
| GAPDH | Mouse | 60004-1-Ig | Protein Tech | 1 in 3000 | WB |
| GAPDH | Rabbit | 10494-1-AP | Protein Tech | 1 in 3000 | WB |
| anti Rabbit IgG NIR 800 | Donkey | A21057 | Licor | 1 in 15000 | WB |
| Anti-Mouse IgG NIR 680 | Goat | 92668071 | Licor | 1 in 15000 | WB |
| Piezo1 | Mouse | NBP2-75617 | Novus Bio | 1 in 1000 | WB |
| VE-Cadherin | Mouse | sc-52751 | SantaCruz | 1 in 10000 | IF |
| PECAM-1 | Goat | sc-1506 | SantaCruz | 1 in 1000 | IF |
| Mouse IgG control |  | AF007 | RandD | see methods | Beads |
| Mouse IgG control |  | MAB002 | RandD | see methods | Beads |

**Table S1:** Antibodies used for in vitro research.

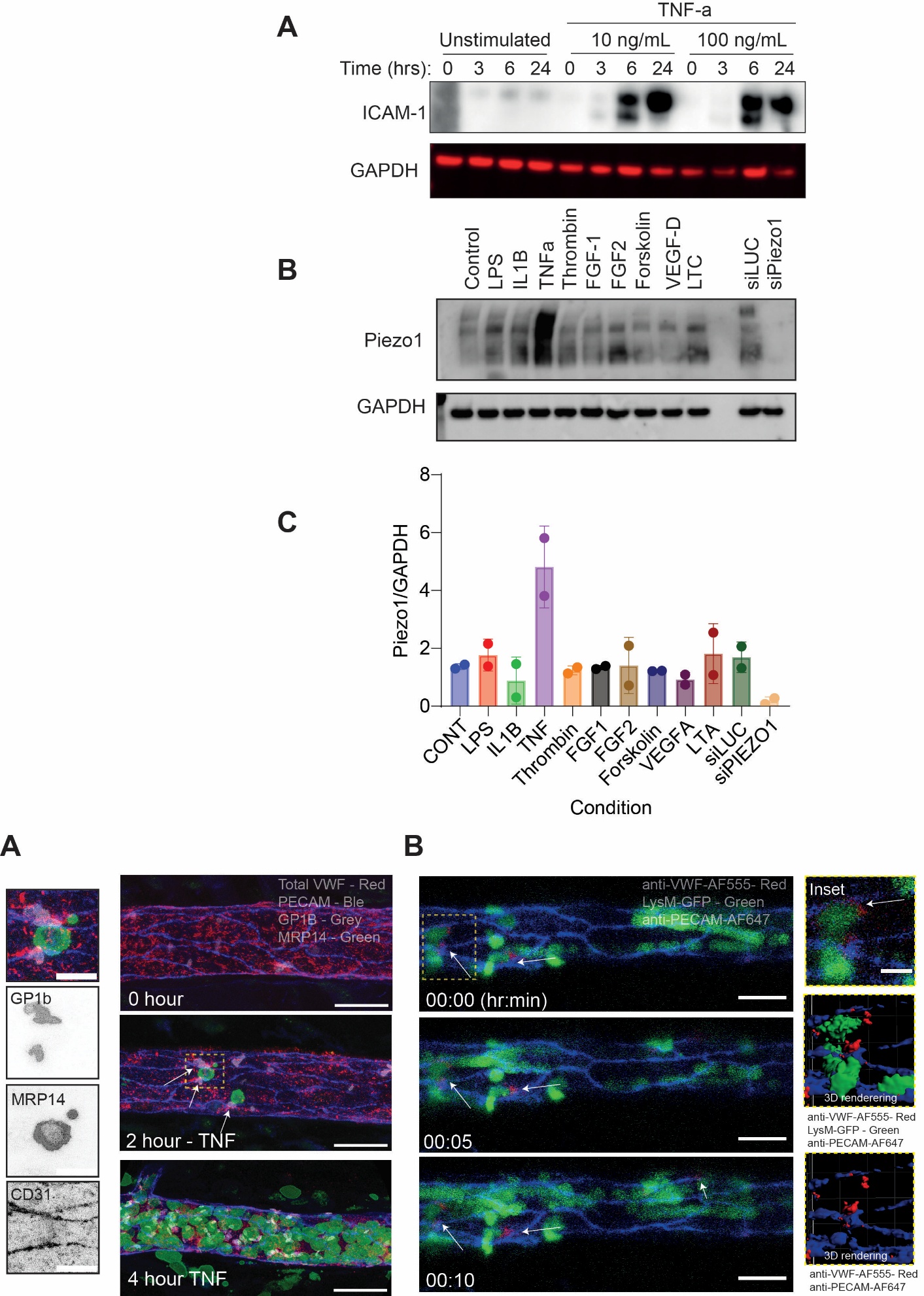

**Figure S1: TNF increases expression of ICAM-1 and Piezo1 at the protein level. (A)** HUVEC were exposed to a range of concentrations (10-100ng/mL) TNF and incubated for a time course of 0,3,6 and 24 hours. Western immunoblotting determined ICAM-1 expression as early as 6 hours post-stimulation. A time point of 16 hours was chosen to for assaying the effect of ICAM-1 binding beads. **(B)** Immunoblotting for Piezo1 determined increased expression of Piezo1 in TNF inflamed HUVEC. Control (luciferase targeting) and Piezo1 targeting siRNA treated cells were used to illustrate specificity of antibody. **(C)** Densitometric analysis of immunoblotting.

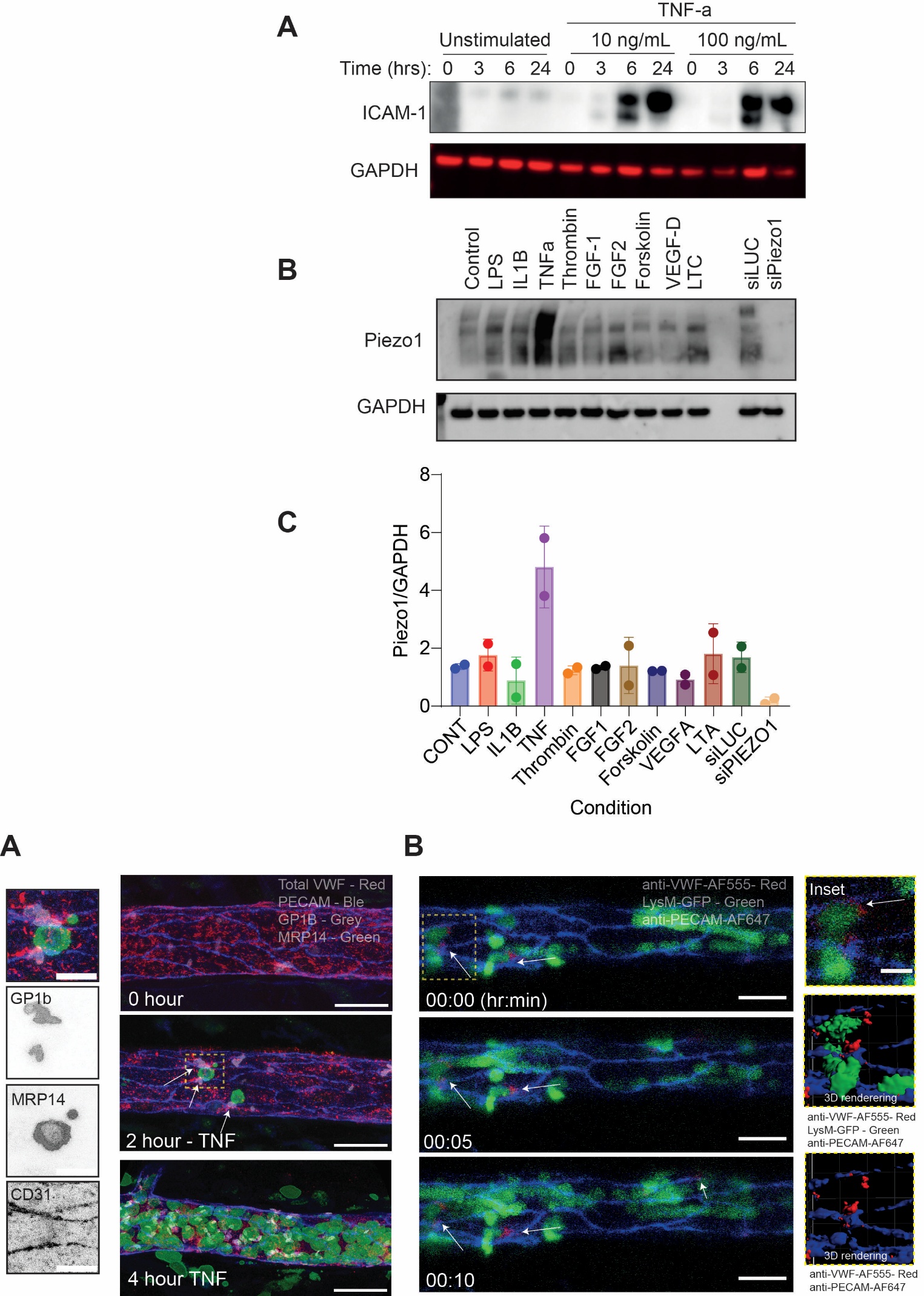

**Figure S2. Extended panels of Figure 2. (A)** Extended time points for Figure 3J, Four-color confocal imaging of wholemount fixed cremaster muscle tissue confirmed platelet adherence at neutrophil trans-endothelial migration (TEM) sites. **(B)** Extended time points for Figure 3K. Anti-VWF-AF555 (5 µg IV) and anti-PECAM-1-AF647 (3 µg intrascrotal) were administered to LysM-GFP mice to visualize VWF secretion during inflammation in the cremaster muscle. Intravital imaging demonstrated VWF secretion at sites of neutrophil adhesion. Arrows indicate adhered neutrophil encapsulated by VWF signal. Scale bar 30 µm.

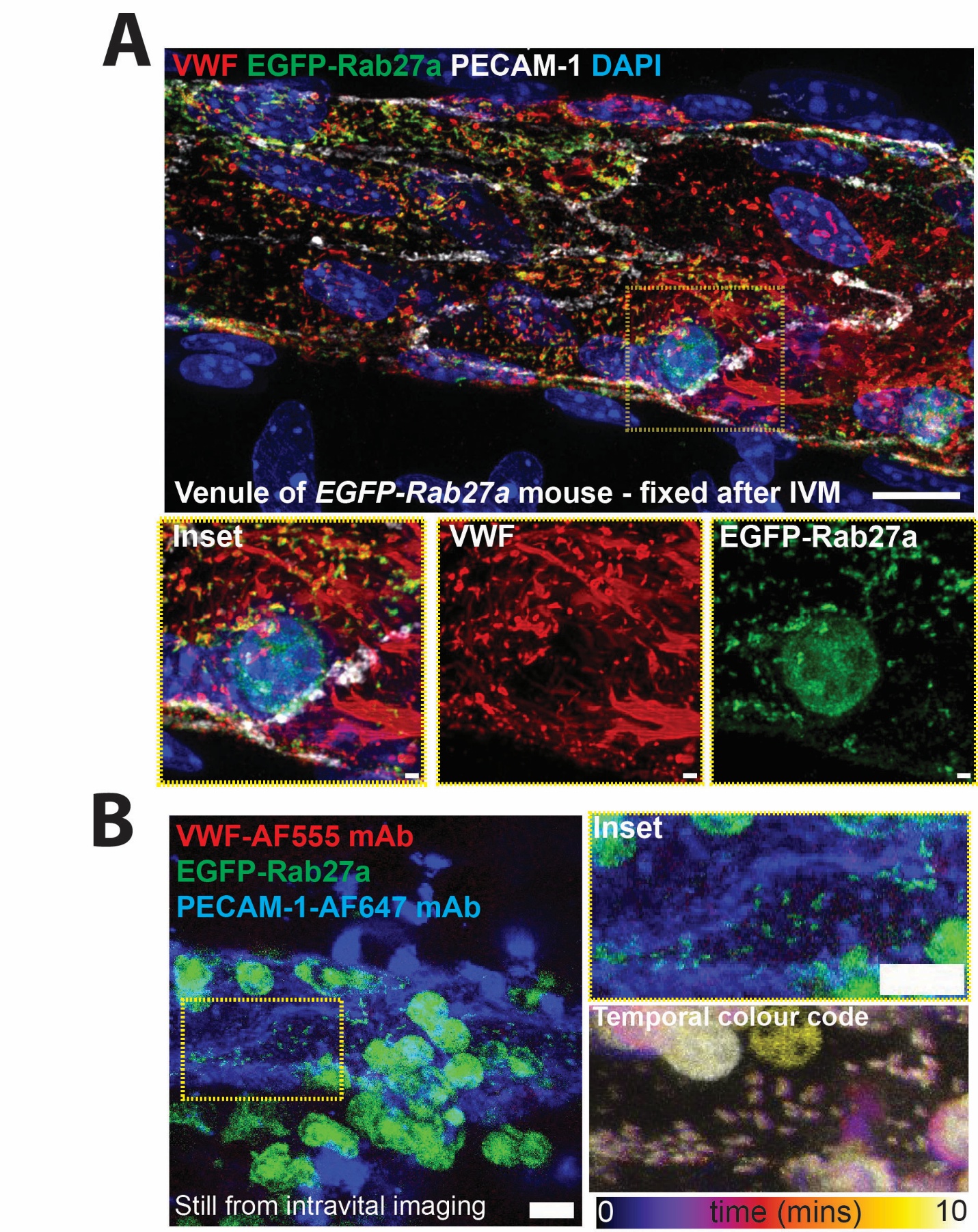

**Figure S3. Confocal microscopy of fixed whole mount cremaster muscle tissue following IVM.** Demonstration of VWF secretion (red) at the site of neutrophil adherence in the *EGFP-Rab27a* mouse. Scale bar 10um. Inset scale bar 1 µm.

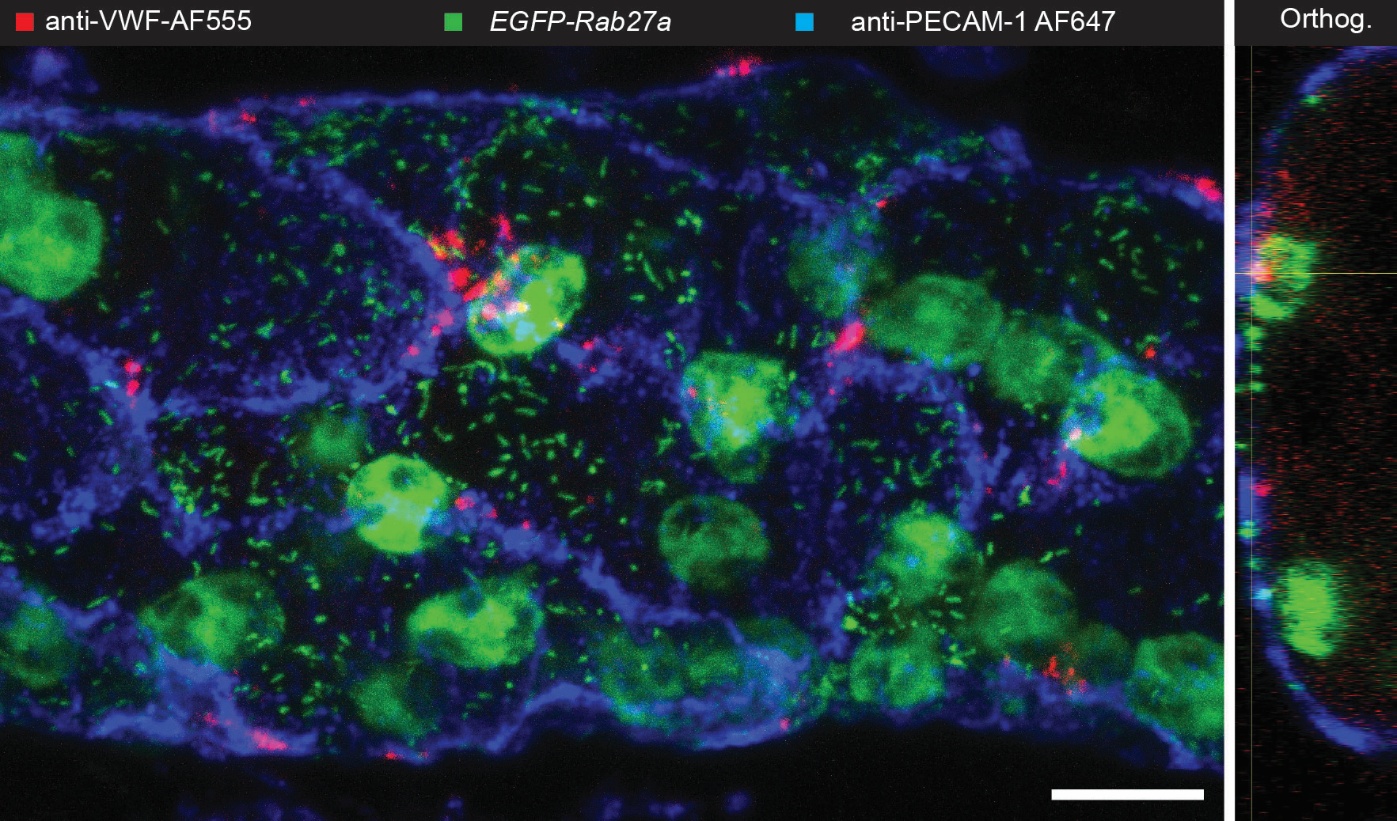
**Figure S4: Spatial profile of VWF secretion in vivo**. Maximum intensity project and orthogonal view of venule from EGFP-Rab27a mouse shows spatial profile and apical secretion of VWF. Histamine stimulation of cremaster muscle. To achieve in situ labelling of secreted VWF, anti-VWF-AF555 was injected intravenously before inflammatory insult. Scale bar 10 µm.
